# Scar-associated endothelial-stellate cellular crosstalk drives fibrosis resolution in MASH

**DOI:** 10.1101/2025.07.08.663293

**Authors:** Kenneth Li, Vardhman Kumar, Tran To, Maylene Yu, Bruno Cogliati, Yashaswini N Chittampalli, Mark Miller, Andrea D Branch, Bruno Giotti, Alexander M Tsankov, Li Chen, Mathieu Petitjean, Chenggang Lu, Yang Li, Sangeeta N Bhatia, Scott L Friedman, Shuang Wang

## Abstract

Fibrosis, or scarring, can affect many organs including liver, lung, heart, kidney, intestines etc. and is responsible for ∼40% of mortality in the industrialized world. Compared to other organs, fibrosis in the liver typically resolves when the source of injury is extinguished. Elucidating the molecular mechanisms that underlie spontaneous fibrosis resolution in the liver may lead to novel antifibrotic strategies for all organs. In this study we established a robust mouse model of fibrosis regression in MASH (Metabolic dysfunction-Associated Steatohepatitis), a highly prevalent chronic liver diseases worldwide, and performed single cell and *in situ* molecular profiling of the liver to define novel drivers of fibrosis regression. As fibrosis regressed, we detected a reduction of inflammatory cells and an expansion of endothelial cells. Prediction of cell-cell communication using the Calligraphy pipeline identified a Wnt9b-Sfrp2 crosstalk that emerges as fibrosis resolved in our model. To establish the Wnt9b-Sfrp2 crosstalk as a driver of fibrosis resolution we treated mice with recombinant Sfrp2, which slowed spontaneous fibrosis regression compared to vehicle treated mice. From our single cell datasets we identified a subset of endothelial cells, termed “Endo4”, as the source of Wnt9b. Immunostaining of the Endo4 marker VWF using tissue clearing and 3D imaging revealed VWF+ vasculature enveloped by activated hepatic stellate cells (HSCs) that penetrated deep into the fibrotic septa, establishing Endo4 as de facto scar-associated endothelial cells and providing a structural basis of their cellular crosstalk with HSCs. Finally, using a recently developed *in situ* protease activity screen, prominent serine protease activity co-localized with both scar-associated Endo4 cells and HSCs. In summary, we uncovered an WNT-dependent endo-stellate crosstalk within the fibrotic niche as a novel regulatory node underlying murine MASH fibrosis regression, and a promising therapeutic target.

## Introduction

Fibrosis, or scarring, underlies ∼40% of deaths in the industrialized world [1]. In the liver, chronic liver disease provokes scar formation (i.e. fibrosis), which can progress to cirrhosis, liver failure, and cancer. An epidemic of metabolic associated steatohepatitis (MASH; previously known as non-alcoholic steatohepatitis (NASH)) threatens the health of at least ∼100 million worldwide [1]. In this disease, fibrosis is the strongest predictor of patient mortality [2]. Despite progress in fibrosis research, for all organs there exists only two modestly effective antifibrotic drugs in lung fibrosis and no approved therapies for liver fibrosis [1]. Addressing this major unmet medical need demands entirely new therapeutic strategies to attack organ fibrosis.

In some organs, most notably liver, fibrosis regresses spontaneously when the source of injury is extinguished. This is exemplified in patients who are cured of chronic hepatitis C virus infection with direct acting antivirals, where even hepatic scar that has accumulated for decades, consisting of fibrillar collagens, spontaneously regresses, while liver function improves [3]. Spontaneous resolution of liver fibrosis has been reproduced in mouse models, including carbon tetrachloride induced chemical injury and several mouse models of MASH [4]. However, the fibrosis field lacks a systematic understanding of how scar resolves in the liver, especially in the context of MASH.

We reason that mechanistic insights into spontaneous fibrosis regression in the liver will lead to novel antifibrotic strategies to treat chronic liver diseases where the source of injury cannot be extinguished, and in other organs where fibrosis is considered irreversible [1]. Although the mechanism remains unclear, several liver cell types expressing a panel of candidate proteases and their inhibitors have been purported to regulate liver fibrosis resolution. Restorative macrophages (defined as Ly-6C^lo^ in mouse) are implicated in fibrosis regression, as their depletion or expansion can delay or speed up fibrosis resolution *in vivo*, respectively [5, 6]. Hepatic stellate cells (HSCs) that activate into the main fibrogenic cells in the liver release a slew of proteases which may contribute to matrix turnover [4, 7]. These studies on macrophages and HSCs have pointed to their expression of specific MMPs and their inhibitors as downstream mediators of extracellular matrix turnover resulting in net fibrosis accumulation or degradation [4, 8, 9]. Other cell types implicated in fibrosis resolution in liver include neutrophils and liver sinusoidal endothelial cells, although their contributions are suggested to be indirect through modulation of macrophages and HSCs, respectively [4, 10]. Together these studies are beginning to hint at the molecular mechanisms underlying spontaneous fibrosis resolution in the liver, which likely engage a complex and exquisitely coordinated response from multiple cell types expressing a range of matrix-degrading proteases that is yet to be defined, especially for MASH fibrosis.

Unprecedented advances in molecular characterization of tissue samples spatially (2D and 3D) and at single cell level are revolutionizing our understanding of complex disease mechanisms. Notably single cell/nucleus and spatial RNA-sequencing simultaneously capture gene expression across different cell types and spatial contexts. When coupled with ingenious computational pipelines, they can be used to interrogate the entire landscape of intercellular crosstalk along with the ligand/receptor pairs involved [11]. Applying one such computational pipeline to our snRNAseq datasets to uncover cell-cell interactions in MASH mice and patients, we previously discovered a complete rewiring of intercellular communication in late-stage MASH fibrosis culminating in the emergence of a stellate cell autocrine signaling circuit [12]. In another line of cutting-edge technological developments, *in situ* zymography techniques directly detect and map protease activity across tissue samples. Application of our Activatable Zymography Probes to tumor samples uncovered unique protein substrates and protease activity within the tumor vasculature, pointing to the possibility of enzymatic activity-based disease detection and therapeutic targets [13, 14]. These technological advances in RNA sequencing and protease activity detection from our laboratories and others set the stage for deconvoluting complex molecular mechanisms driving spontaneous fibrosis regression in MASH while elucidating novel antifibrotic strategies and drug targets.

In this study we applied cutting-edge single cell/spatial RNA sequencing, tissue-clearing/3D imaging, and *in situ* protease activity detection to a temporally defined mouse model of spontaneous MASH fibrosis resolution. This combinatorial approach allowed us to deconvolute the complex molecular niche surrounding the hepatic scar, leading to the identification of scar-associated HSC and endothelial cell subsets exhibiting serine protease activity that crosstalk extensively through WNT-SFRP to drive fibrosis resolution.

## Results

### Characteristics of fibrosis regression in a murine MASH model

To establish a robust MASH fibrosis regression model, we first placed mice for 6 weeks on our previously established FAT-MASH murine MASH model [15]. We found that fibrosis in this model significantly resolved at 2 weeks and further reduced at 6 weeks following returning to control conditions and injury cessation (MASH Regression, **Figure 1**). As a comparison, we also ran a chronic CCl_4_ model for 6 weeks and measured fibrosis regression at 2 and 6 weeks following fibrosis resolution (CCl_4_ Regression, **Figure 1A**). Average fibrosis levels as measured by Picosirius red staining of collagen decreased at both 2wk and 6wk Regression timepoints in both models compared to Peak Fibrosis. The collagen positive area in the 2wk and 6wk Regression timepoints were decreased by an average of 11% and 46%, respectively, in the MASH Regression model, and 34% and 65% in the CCl_4_ Regression model, indicating a consistent and rapid decrease across both models (**Figure 1B**,**C**). Reduced fibrosis levels were accompanied by visible reductions in the histological hallmarks of MASH, including steatosis, inflammation, and hepatocyte ballooning (**Figure S1**).

**Figure 1.**
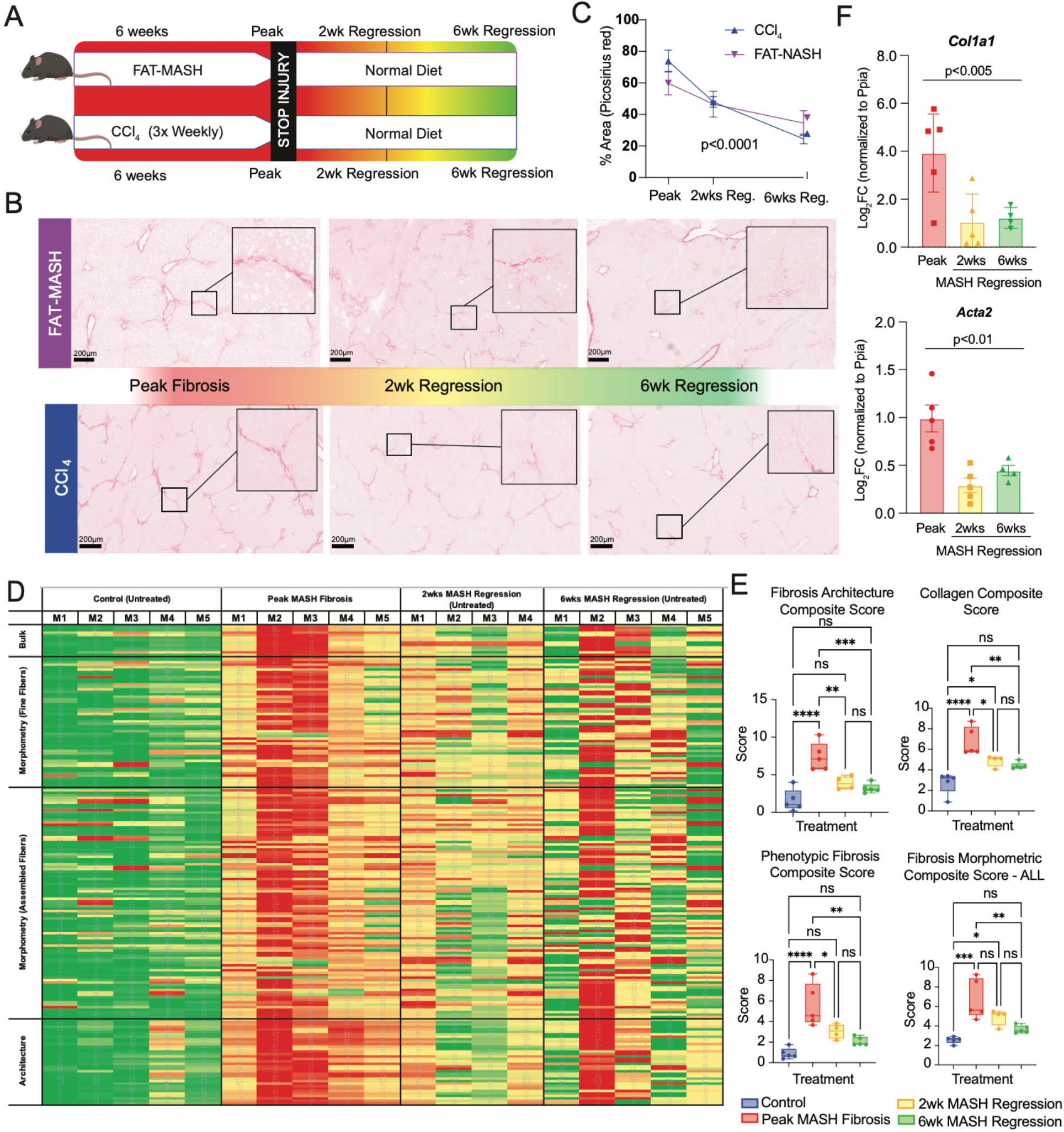
A mouse model for studying spontaneous fibrosis resolution in MASH. (**A**) Schematic of experimental layout. Fibrosis was induced in mice with either FAT-MASH or CCl4 regimen for 6 weeks (peak fibrosis) then returned to control conditions to allow fibrosis regression for 2 weeks (2wk regression) or 6 weeks (6wk regression). (**B**) Collagen positive area for Peak, 2wk Regression, and 6wk Regression samples as determined by Picrosirius red staining. Representative images were taken with 5X objective. (**C**) Quantification of collagen positive area for each mouse, as a percent of the corresponding model’s Peak Fibrosis collagen positive area. N=6-8 for CCl_4_ Peak, 2wk, and 6wk Regression timepoints, and N=12-14 for MASH Peak, 2wk, and 6wk Regression timepoints pooled across multiple studies. (**D**) Digital pathology analysis of mice from a single experiment reveals changes in liver tissue architecture during fibrosis regression. Each mouse is represented by the column. Each row is a parameter measured by the Fibronest platform. Heatmap colors (red=high, green=low) are standardized across each parameter. N=4-5 for MASH Peak, 2wk, and 6wk Regression timepoints. (**E**) Composite fibrosis scores calculated on the basis of (D). (**F**) Gene expression of *col1a1* and *acta2* quantified via qPCR. N=4-5 for Peak, 2wk regression, 6wk regression, and untreated timepoints.

To extensively characterizes both absolute collagen content and important statistical features of the distributions of collagen fibers’ morphometric and architectural phenotypes. across various MASH regression timepoints, we applied the Fibronest digital pathology platform to whole-slide scans of Picosirius red staining from mice from a single MASH regression study. As expected, we detect a significant reduction in fibrosis severity at both 2 and 6wk MASH Regression timepoints compared to peak fibrosis, however, no significant difference was detected between 2wk and 6wk MASH Regression (**Figure 1D**,**E**). This was corroborated by measurement of fibrogenic gene expression (*Col1a1* and *Acta2*) in liver homogenates by qPCR, demonstrated significant reduction at 2wk MASH regression that is maintained at 6wk MASH regression (**Figure 1F**). The alterations during regression in these key readouts of fibrosis in the MASH Regression model demonstrate that this model and timepoints are suitable to study liver fibrosis regression.

### Transcriptomic landscape of spontaneous fibrosis resolution at single-cell resolution

To obtain a holistic view of transcriptomic changes that accompany fibrosis resolution in mice, we first performed single cell RNA sequencing (scRNAseq) in both CCl_4_ and MASH regression models at Peak Fibrosis, 2wk, and 6wk Regression timepoints (**Figure 2A**). Informatically merging datasets from Peak Fibrosis, 2wk, and 6wk Regression timepoints from both models and clustering using a Uniform Manifold Approximation and Projection (UMAP)– based approach revealed the predominant presence of immune cells and endothelial cells (**Figure 2B**). Specifically, B-cells and endothelial cells comprised the highest (26%) and second highest (24%) proportion of cells detected (**Figure S2A**). Top 2 genes markers expressed by each cell subset detected are shown in **Figure 2C**. Interrogation of changes in cell type abundance during regression in the scRNAseq dataset revealed dynamic changes in the immune and endothelial cell compartment. Compared to Peak Fibrosis, B cells decreased by 26% and 19% during 6 weeks of CCl_4_ and MASH Regression, respectively, likely reflecting reduced inflammation following injury cessation (**Figure 2D**). On the other hand, endothelial cells increased by 13% and 21% during 6 weeks of CCl_4_ and MASH Regression, suggesting active repair of the tissue microenvironment (**Figure 2D**).

**Figure 2.**
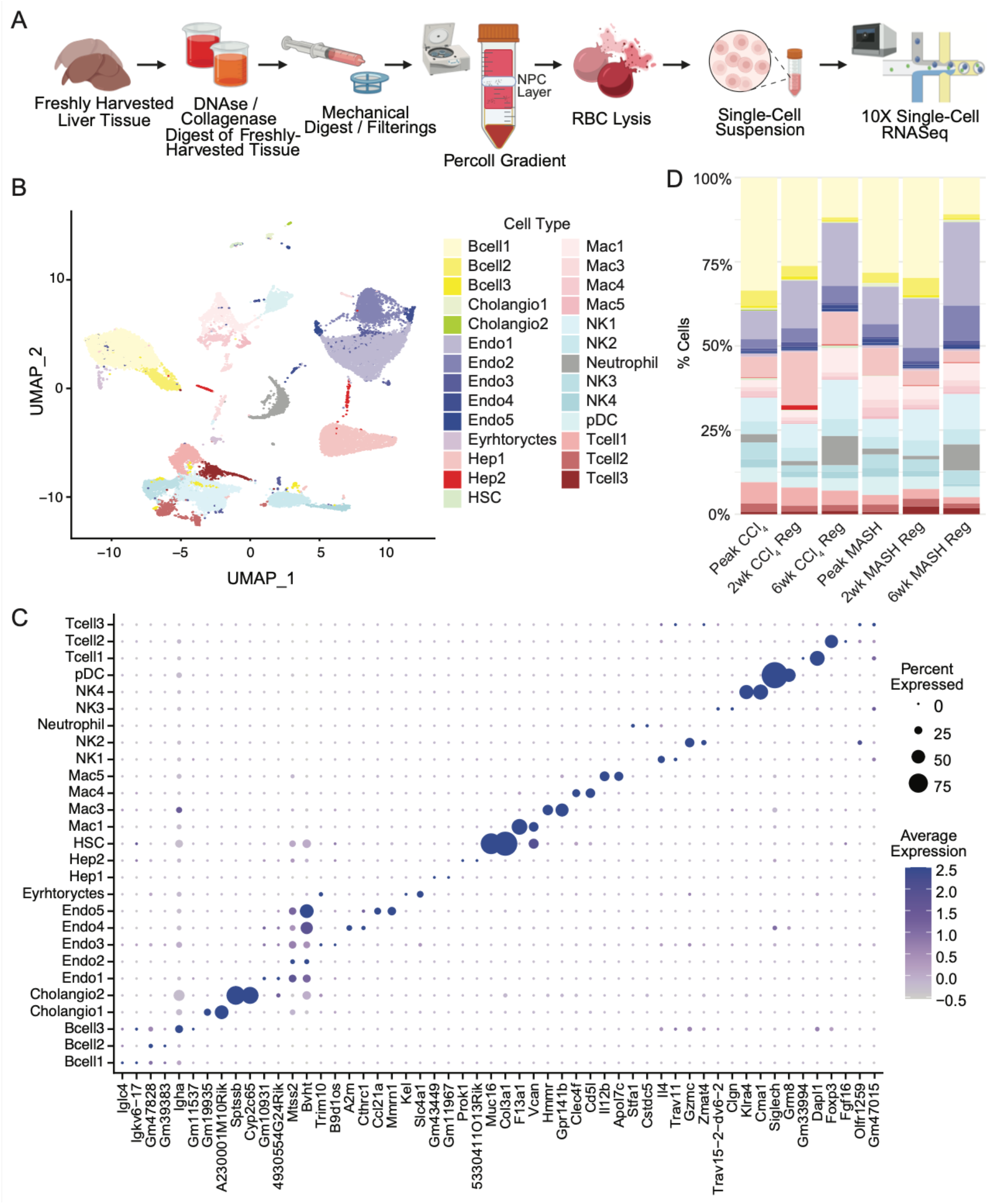
Transcriptomic landscape of fibrosis regression by single cell RNA-seq. (**A**) Schematic of scRNAseq experimental layout from cell isolation to sequencing. (**B**) UMAP visualization of cell clusters colored by cell type. “Cholangio” = cholangiocytes, “Endo” = endothelial cells, “Hep” = hepatocytes, “HSC” = hepatic stellate cells, “Mac” = macrophages, “NK” = natural killer cells, “pDC” = Plasmacytoid dendritic cells. (**C**) Top two marker gene expression for each cell type (by log2 fold change). (**D**) The proportion of annotated cell types at each time point, relative to the total number of cells in each time point.

Previous studies from our laboratory and others have demonstrated differential cell capture between single nucleus and single cell RNAseq. To comprehensively profile the transcriptomic landscape of the liver undergoing fibrosis resolution, we additionally performed single nucleus RNA-seq from Peak Fibrosis, 2wk, and 6wk Regression timepoints of both the CCl_4_ and MASH models (**Figure 3A**). UMAP-based clustering analysis of the merged snRNAseq data from Peak Fibrosis, 2wk, and 6wk Regression timepoints from both MASH and CCl_4_ models revealed the most abundant cell type being hepatocytes, comprising 68% of all cells, followed by endothelial cells (16%) and HSCs (8%) (**Figure 3B, C, Figure S2B**), in line with previous studies including from our laboratory showing snRNAseq capture liver cell types at their expected proportions [12, 16]. Interrogation of cell type abundance changes during regression in the snRNAseq dataset revealed a consistent expansion of the endothelial cell compartment at 2wks Regression for both CCl_4_ and MASH models (**Figure 3D**). Furthermore, we detect a cluster of vimentin-expressing endothelial cells (“Endo5”) at Peak Fibrosis and 2wks Regression timepoints that disappeared at the 6wk Regression timepoint in both models, which may undergo endothelial-mesenchymal transition (**Figure S3**).

**Figure 3.**
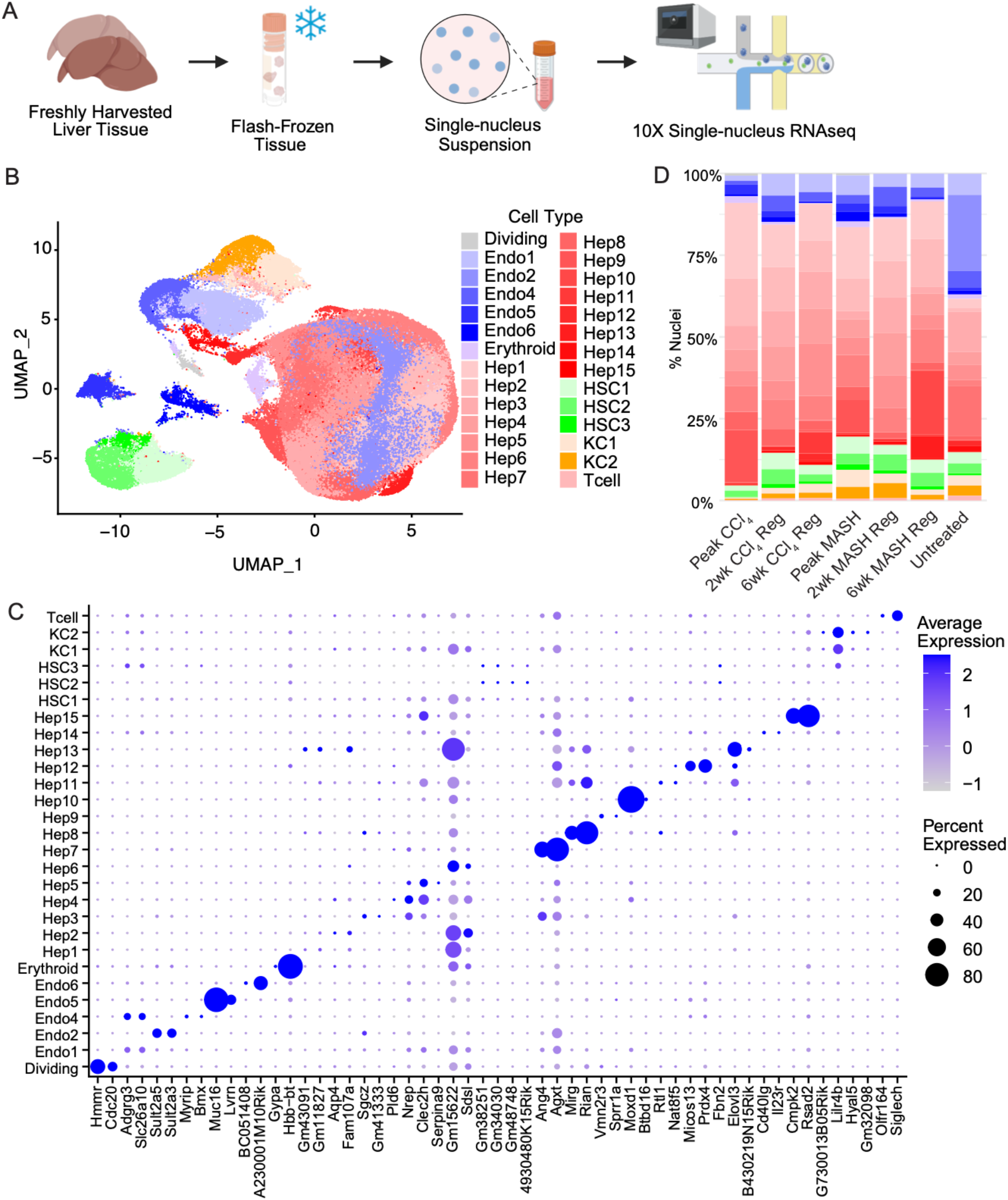
Transcriptomic landscape of fibrosis regression by single nucleus RNA-seq. (**A**) Schematic of snRNAseq experimental layout, from nucleus isolation from flash frozen liver tissue to sequencing. (**B**) UMAP visualization of cell clusters colored by cell type. “KC” = Kupffer cells, “HSC” = hepatic stellate cells, “Hep” = hepatocytes, “Endo” = endothelial cells. (**C**) Top two marker gene expression for each cell type (by log2 fold change). (**D**) The proportion of annotated cell types at each time point, relative to the total number of cells in each time point.

### A WNT-SFRP crosstalk drives fibrosis resolution in MASH

To study the cell-cell crosstalk underlying fibrosis regression, we performed CellphoneDB analysis [17] that predict receptor-ligand interactions based on their expression on annotated cell types in the snRNAseq datasets of the CCl_4_ and MASH Regression models. In line with studies from our laboratories and others, we again detect highest number of interactions between hepatic stellate and endothelial cells [12, 18]. Mirroring our previous report of increased cell-cell communication in MASH [12], we detect a global decrease in intercellular signaling, represented by the total number of significant interactions predicted across all cell types, as fibrosis regressed in both MASH and CCl_4_ models (**Figure S4**). Despite a global decrease in cell-cell communication, we found that interactions between hepatic stellate and endothelial cells remain to be the most prominent as fibrosis resolves (**Figure S4**).

Although CellphoneDB analysis provided a global view of the cell-cell communication landscape associated with fibrosis regression, we recovered hundreds of significant interactions making it difficult to prioritize hits for functional validation. To further prioritize interaction pairs that are most biologically relevant, we applied the recently published Calligraphy pipeline [19] and its modular-based signal discovery methods to our single-nucleus and single-cell sequencing datasets (**Figure 4A**). Using a co-expression modular analysis, Calligraphy analysis identified much fewer significant interactions at Peak Fibrosis, 2wk, and 6wk Regression timepoints (**Figure 4B**, complete list in **Table S1**). Applying Calligraphy to the integrated CCl_4_ and MASH Regression datasets, we identified a distinct interaction network between endothelial cells, HSCs, and cholangiocytes, along with 3 other interaction networks between immune cells (neutrophils/NK/BCell) and hepatocytes. Focusing on the interaction network between endothelial cells, HSCs, and cholangiocytes that is present at Peak Fibrosis and also persist at 2wks Regression, a period of maximum fibrosis resolution in our MASH Regression model identified in **Figure 1**, we found that this interaction network centers around WNT signaling engaging both WNT ligands (Wnt7a/b, Wnt9b) and inhibitors (Sfrp1/2) (**Figure 4B**).

**Figure 4.**
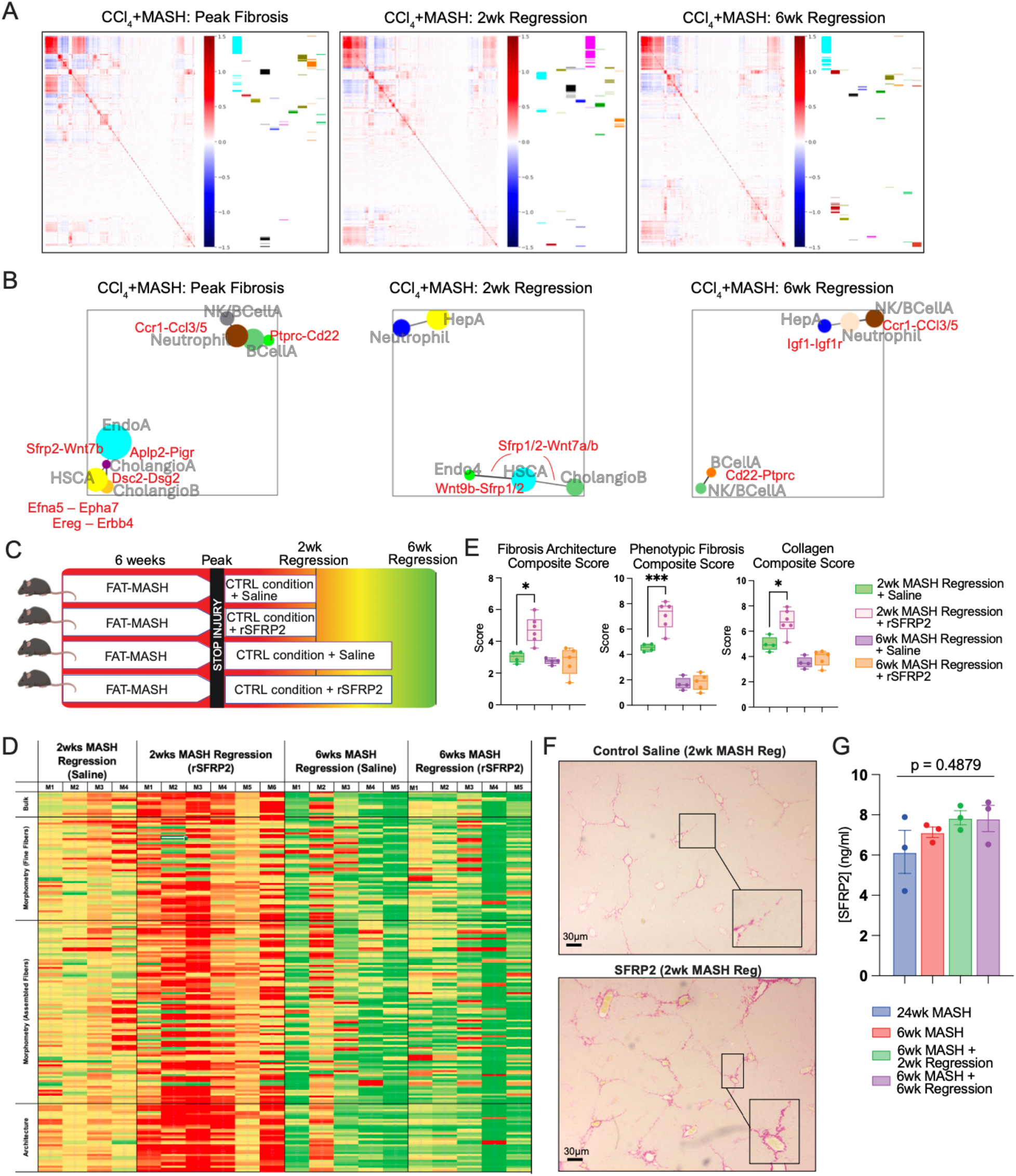
Fibrosis resolution in MASH is mediated by a WNT-SFRP crosstalk between hepatic stellate and endothelial cells. (**A**) Heatmaps showing the correlation among the combined receptors and ligands with each other in the different timepoints of the scRNAseq dataset as predicted by Calligraphy [19]. Right panels show the modules of co-regulated gene activity derived from the correlation heatmap. (**B**) Modular interaction networks predicted by Calligraphy based on the modules of co-regulated gene activity at each time point. Annotations show the annotated cell type(s) that likely contribute to the module. Mediators of module-module interactions that are predicted to be significant by Calligraphy are shown in red, with complete list provided in Table S1. (**C**) Schematic of experimental design. C57BL6/J mice were fed FAT-MASH diet for 6 weeks to induce fibrosis, then switched back to control conditions for regression. Mice were regressed for 2 or 6 weeks, during which they received either twice-weekly intraperitoneal injections of SFRP2 or saline vehicle. (**D**) Digital pathology analysis revealed changes in tissue architecture during regression with or without SFRP2 treatment. Each column is a mouse. Each row is a parameter measured by the Fibronest platform. Heatmap colors (red=high, green=low) are standardized across each parameter. (**E**) Composite fibrosis scores calculated on the basis of (D). (**F**) Representative collagen positive area for 2wk MASH Regression mice treated with SFRP2 or saline vehicle, as shown by picrosirius red staining. Images were taken at 5X objective. (**G**) Serum SFRP2 levels in mice treated under MASH model for 6 weeks and regressed for either 2 or 6 weeks as measured by ELISA, also including mice on MASH model for 24 weeks as control. N=3 for all timepoints.

WNT signaling coordinates tissue repair across multiple organs. Recent studies have shown that pericentral endothelial cells provide the crucial WNT ligands (Wnt9b, Wnt2) that establishes the metabolic zonation of hepatocytes and promote their proliferation in liver regeneration [20, 21]. To establish the functional relevance of WNT-SFRP crosstalk in MASH fibrosis regression we systemically injected recombinant SFRP2 twice weekly for 2 or 6 weeks following injury cessation in the MASH Regression model and tested its effect on fibrosis regression (**Figure 4C**). In this study, two weeks of SFRP2 treatment during fibrosis regression blocked spontaneous fibrosis resolution in these mice as measured by area and AI-based quantification of Picrosirius red staining (**Figure 4D-F**). However, we found that the effect of SFRP2 treatment disappears by 6 weeks of fibrosis regression, suggesting that recombinant SFRP2 cannot exert a permanent brake on fibrosis resolution (**Figure 4D,E**). Finally, SFRP2 is clearly detectable in the serum of both MASH mice and DAA-treated HCV patients undergoing fibrosis resolution using ELISA with no significant directional change with disease status (**Figure 4G, Figure S5**).

### A subset of Wnt9b+ Vwf+ Endothelial Cells Vascularize Within Fibrotic Scar

To better understand how this WNT-SFRP crosstalk could contribute to fibrosis resolution, we first searched for the cellular source of Wnt9b. We found that Wnt9b is most highly expressed by a subset of endothelial cells we refer to as “Endo4” in our snRNAseq datasets, which is also marked by expression of Vwf and Ace (**Figure 5A,B**). During fibrosis regression of both MASH and CCl_4_ models, the highest proportion of Endo4 is found at the 2wks Regression timepoint (**Figure 5C**). On spatial RNA-seq data from Peak MASH Fibrosis, we found a colocalization of Endo4 markers with the fibrotic septa (marked by white arrows) along with marker of activated HSCs (Acta2), suggesting that Endo4 may represent a subset of scar-associated endothelial cells (**Figure 5D**). To conclusively establish Endo4 as scar-associated endothelial cells, we carried out tissue-clearing and 3D imaging of the endo4 marker VWF in livers of MASH mice at Peak Fibrosis and 2wk Regression using the iDISCO pipeline [22]. In these mice, HSCs are genetically labelled with a TdTomato fluorescent reporter activated by the Cre recombinase driven by the *Lrat* promoter [23]. At Peak Fibrosis, we found VWF+ endothelial cells not only strongly co-localize with the fibrotic septa marked by TdT+ HSCs with flat, sheet-like myofibroblast morphology, they actually formed vascular structures within these fibrotic regions and these structures persisted at 2wks Regression even as HSCs begin to regain their quiescent, neuronal morphology and pull away from the scar regions (**Figure 5E**).

**Figure 5.**
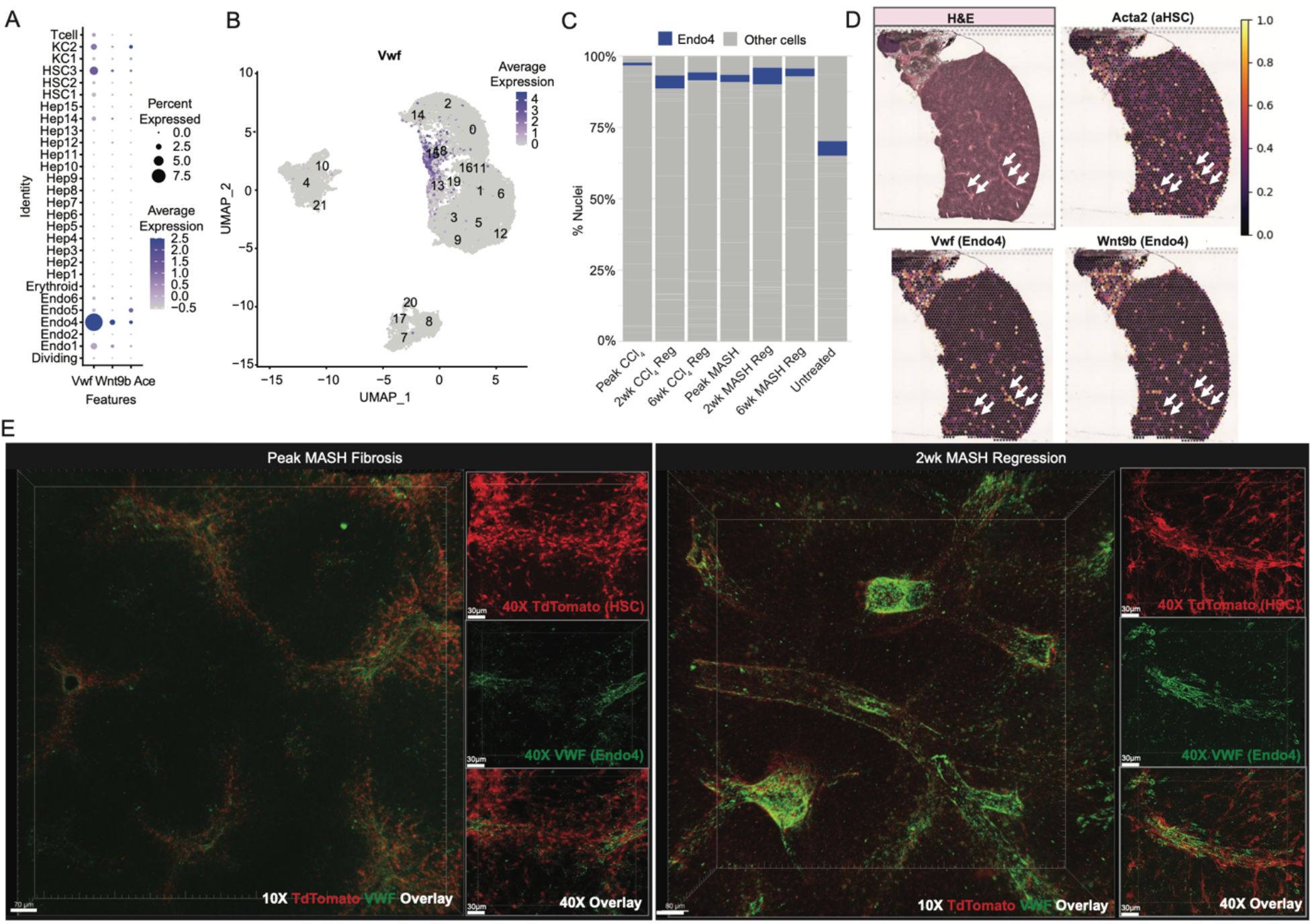
A subset of VWF+ WNT9b+ endothelial cells vascularize within the fibrotic scar. (**A**) Dot plot showing percent and average expressions of *Vwf, Wnt9b*, and *Ace* in snRNAseq clusters. (**B**) UMAP embedding of the endothelial cell subsets from the integrated CCl_4_ and MASH Regression snRNAseq dataset, color scale indicate expression of Endo4 marker Vwf. (**C**) Percent abundance of Endo4 (red highlighted) subcluster at the different time points in the CCl_4_ and MASH Regression snRNAseq datasets. (**D**) Deconvolved spatial transcriptomic enrichment of *Acta2, Vwf*, and *Wnt9b* at MASH Peak Fibrosis overlayed with hematoxylin & eosin staining. (**E**) TdTomato and VWF protein expression in iDISCO cleared and antibody-stained livers from mice with TdTomato+ HSCs at Peak MASH Fibrosis and 2wk MASH Regression timepoints imaged with 10X objective or 40X objective.

### Scar-associated hepatic stellate and endothelial cells exhibit serine protease activity

To search for proteases that degrade the extracellular matrix in our MASH regression model, we utilized a global database of proteases and inhibitors and analyzed their scaled expression in our MASH and CCl_4_ Regression datasets. In addition to macrophages, which has shown to produce ECM remodeling MMPs, we also found high expression of proteases in endothelial cells and HSCs in both scRNAseq and snRNAseq datasets (**Figure S6**). Among endothelial cell subsets, the previously described Endo4 exhibited high expression of proteases, including matrix metalloproteases and serine proteases, that is distinct from other endothelial cell subsets. Specifically, Endo4 upregulates an ECM-remodeling signature enriched for collagenases and gelatinases along with lysosomal cathepsins. HSCs expressed a matrix-remodeling module as well as an immune-related module with elements of complement activation, which is consistent with their known role in orchestrating immune recruitment during scar clearance. Different subtypes of endothelial cells contribute proteolytic enzymes during regression, positioning endothelial cells as one of the major matrix breakdown cell types during regression in addition to HSCs and macrophages.

Ultimately proteases are defined by their enzymatic activity. To identify proteases and their cellular sources that contribute to ECM turnover in our MASH Regression model we screened Activable Zymography Probes (AZP) on liver cryosections from Peak Fibrosis as described previously ([13, 14], **Figure 6A**). From this screen we identified AZP1 as a top hit which demonstrated strongest signal at Peak Fibrosis and reduced signal at 2wks Regression (**Figure 6B,C**). To identify the cellular source of AZP1 signal we also co-localized AZP1 signal with makers of HSCs (DES), Endo4 (VWF), and cholangiocytes (CK7). We found strong co-localization between AZP1 and HSCs followed by Endo4, both reaching levels comparable to colocalization with the collagen positive control (**Figure 6D,E**). On the other hand much lower co-localization was found between AZP1 with cholangiocyte maker CK7 (**Figure 6D,E**). To narrow down on the class of protease activity being detected by AZP1 we co-treated tissue samples with protease inhibitors against all proteases (broad spectrum), MMPs (Marimastat), and serine proteases (AEBSF). We found that co-treatment with serine protease but not MMP inhibitors blocked AZP1 signal in addition to broad spectrum protease inhibitor (**Figure 6F,G**). These results suggest that serine proteases secreted by scar-associated HSCs and endothelial cells could actively contribute to scar degradation in MASH fibrosis.

**Figure 6.**
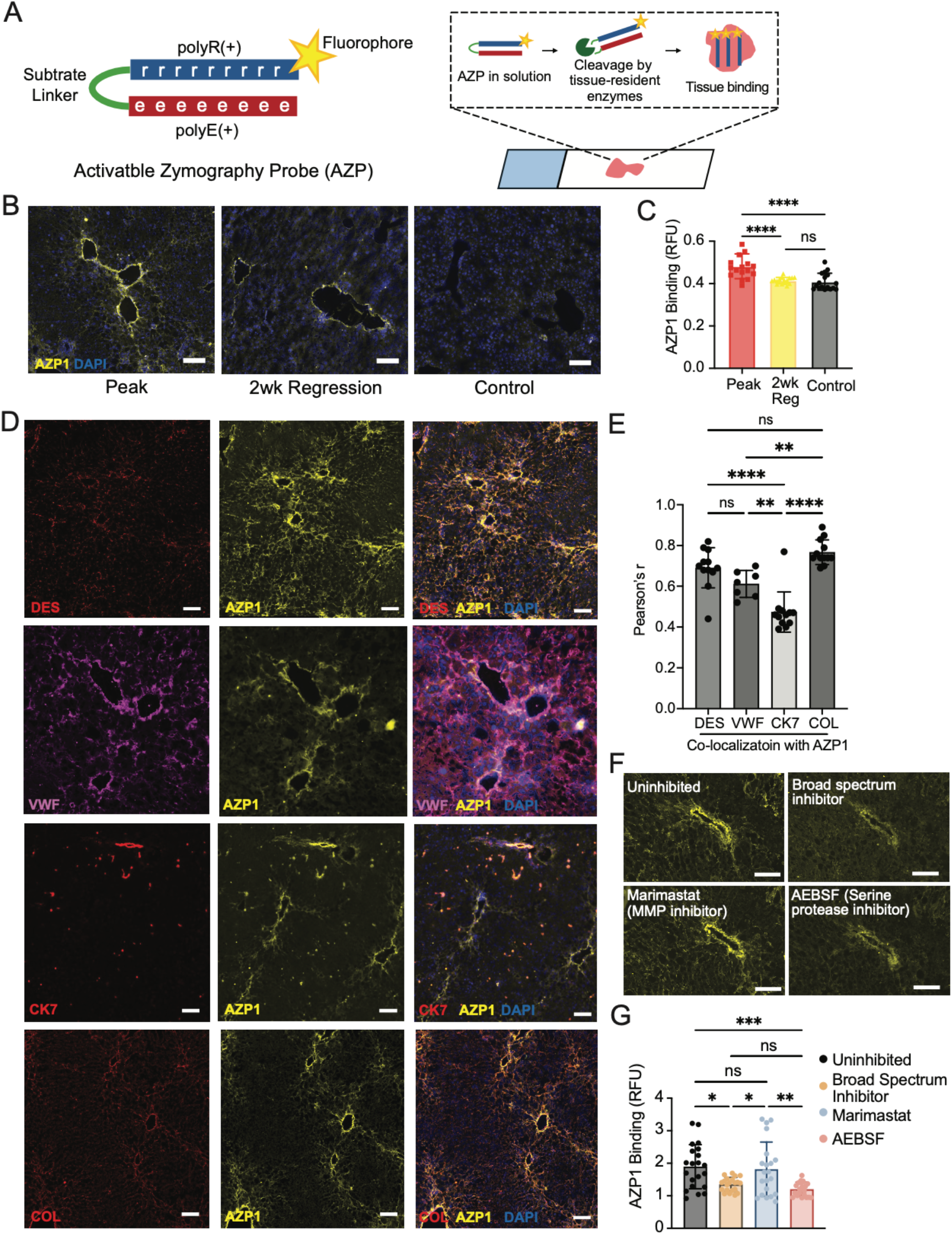
Scar-associated hepatic stellate and endothelial cells exhibit serine protease activity. (**A**) AZP schematic and working principle. (**B**) AZP staining in MASH Peak Fibrosis, 2wk Regression and Control samples and (**C**) quantification. (**D**) AZP stain co-localizes with Desmin, VWF, and Collagen Type I and (**E**) co-localization quantification. (**F**) AZP staining in the presence of broad-spectrum inhibitor, marimastat (metalloprotease inhibitor) and AEBSF (serine protease inhibitor) and (**G**) quantification.

## Discussion

In this study we first established and characterized a rapid and robust murine model of spontaneous MASH fibrosis resolution. Using single cell/spatial transcriptomics and protease activity profiling to dissect the complex cellular and molecular interactions that underlie spontaneous fibrosis resolution in our model, we identified a cellular crosstalk between scar-associated HSCs and endothelial cells, mediated by Wnt9b-Sfrp2, that drive fibrosis resolution possibly through secretion of serine proteases.

Unlike fibrosis of other organs, fibrosis in the liver rapidly resolves following injury termination such as following successful HCV clearance. The molecular mechanism behind spontaneous fibrosis resolution in the liver has remained elusive and likely involve intricate coordination between multiple cell types that secrete proteases to degrade the extracellular matrix that make up the hepatic scar. To capture this cellular dynamics in our model and maximize cell recovery, we performed temporal single cell and single nucleus RNAseq during fibrosis regression following both MASH and CCl_4_ injury. We found similar changes in cell composition as fibrosis resolves in both injury models, with a reduction of infiltrating inflammatory cells and a notable expansion of endothelial cells that has not been reported previously. Using Calligraphy to predict significant cell-cell interactions during fibrosis resolution we identified a Wnt9b-Sfrp2 crosstalk at timepoints that correspond to maximum fibrosis resolution. To functionally validate the importance of this Wnt9b-Sfrp2 crosstalk in fibrosis resolution, we injected mice with recombinant Sfrp2 following injury cessation and found that significantly slowed fibrosis resolution, suggesting that systemically manipulating WNT activity could serve as novel antifibrotic strategy in MASH patients. In our datasets Wnt9b is expressed by a subset of endothelial cells that we refer to as “Endo4”, which in the control liver expresses makers of vascular endothelial cells that surround the central veins. In fibrosis, however, we found these cells penetrate into the hepatic scar possibly in response to angiogenic cues due to hypoxia from these regions and form vascular structures that are wrapped by a sheet of activated HSCs. These observations support the development of novel drug delivery strategies to specifically target this fibrotic niche, such as lipid nanoparticles coated with antibodies against the Endo4 marker VWF, which we are currently testing in our laboratory. Developing drugs to specifically target the fibrotic niche may circumvent off-target toxicity that represents a major bottleneck with current antifibrotic strategies such as systemic TGF-β1 inhibition [24].

Finally a screen of protease activity using *in situ* activatable zymography probes identified prominent serine protease activity in the scar regions in proximity to both HSCs and Endo4. To our knowledge the search for proteases that degrade the extracellular matrix in liver fibrosis have mostly focused on MMPs and their inhibitors. Our detection of significant serine protease activity in these scar regions support future investigations into members of the serine protease family as potential therapeutic targets in MASH fibrosis. Furthermore, strongest protease activity was detected at Peak Fibrosis compared to Regression timepoints, which supports a paradigm of ongoing extracellular matrix remodeling during chronic liver injury with the degree of fibrosis reflecting the net result of competing extracellular matrix deposition and degradation processes.

There are several limitations of this study. Although we decided to focus on HSCs, endothelial cells, and their crosstalk from our single cell transcriptomic profiling, there are likely other cellular/molecular players that are engaged in some aspect of fibrosis resolution that we expect to emerge from our datasets as we make our datasets publicly available for the research community to generate novel hypotheses. Systemic administration of SFRP2 may lead to unintended biological effects on other cell types within the liver and on other organs that indirectly affected fibrosis levels in our MASH regression model. In future studies we will search for genetic tools to more specifically manipulate Wnt9b/Sfrp2 levels in the scar-associated endothelial cells or HSCs to conclusively establish their roles in fibrosis resolution. It is also unclear which cells are WNT responsive within our model, which we suspect are scar-associated Endo4 and HSCs as they exhibit highest protease activity in our *in situ* protease activity assays, but remains to be conclusively proven.

## Materials and Methods

### qRT-PCR

RNA was isolated from snap-frozen liver tissue using RNeasy Mini Kit (Qiagen, 74106). Concentration was assessed using an Epoch Biotek microplate reader (Biotek). 1μg of RNA from each sample was reverse-transcribed (42 °C/60 minutes, 70 °C/10 minutes, and 4 °C hold) into complementary DNA using a RNA to cDNA EcoDry Premix (Double Primed) Kit (Takara, 639549). cDNA was diluted (final concentration 4ng/μl in distilled H2O). qPCR was conducted in triplicate with final concentration of 2 ng/μL cDNA from each sample, 1 μM of primers, and 1X SYBR Green Master Mix. Target gene expression was normalized to endogenous levels of *Ppia* or *Polr2a* mRNA (ΔCT) and then displayed as fold change compared to the control group (ΔΔCT).

### Animal Experiments

6-week-old C57B/6J male mice (Jackson) were maintained in ventilated cages (up to 5 mice per cage) in a Helicobacter-free facility on a 12-hour light/dark cycle. Mice were kept on regular chow diet and water for a one-week acclimatization period prior to fibrosis induction. All procedures were approved by the Animal Care and Use Committee of the Icahn School of Medicine at Mount Sinai (IACUC-2018-0063).

### Carbon Tetrachloride (CCl_4_)-only Regression Model

To induce fibrosis, 6-week-old C57B/6J male mice (Jackson) acclimatized at Mount Sinai for 1 week were treated under a well-established fibrosis induction model [25]. Mice were administered with CCl_4_ (Sigma, 289116) dissolved in corn oil by IP injection (0.5μl/g body weight, three times weekly) for six weeks. Mice were fed regular chow diet and tap water, *ad libitum*, during this period. A baseline cohort of “Peak Fibrosis” mice were euthanized 48 hours after the final injection to assess maximum fibrosis response. In separate groups, mice were allowed to recover without treatment (i.e. stop CCl_4_ injections) for up to two (“2wk Regression”) or six (“6wk Regression”) additional weeks.

### MASH Regression Model

To induce MASH, 6-week-old C57B/6J male mice (from Jackson Laboratories) were treated under a well-established FAT-MASH induction model [15]. Mice were administered with CCl_4_ (Sigma, 289116) dissolved in corn oil by IP injection (0.2ul/g body weight, once weekly) for six weeks. Mice were fed a high-fat, high-fructose Western Diet *ad libitum* containing 21.1% fat, 41% sucrose, and 1.25% cholesterol by weight (Teklad, TD.120528) and a high sugar solution containing 23.1g/L D-Fructose (Sigma-Aldrich, F0127) and 18.9g/L F-Glucose (Sigma-Aldrich, G8270) diluted in autoclaved water during fibrosis induction. A baseline cohort of “Peak Fibrosis” mice were euthanized 48 hours after the final injection to assess maximum fibrosis response. In separate groups, mice were allowed to recover without treatment (i.e. returned to normal chow and water and stopped CCl_4_ injections) for up to two (“2wk Regression”) or six (“6wk Regression”) additional weeks.

### Tissue Histology

Fresh liver tissue was fixed overnight in 10% buffered formalin (Epredia #22-050-105) and proceeded to paraffin embedding into paraffin blocks. To make FFPE slides, blocks were sectioned by microtome and mounted onto charged glass slides. For immunostainings, slides were deparaffinized using Xylene (Fisher HC700) and rehydrated with an ethanol gradient (Decon laboratories #2705, 100%/100%/95%/85%/70% ethanol in water). Deparaffinized slides were then boiled in 10mM Citrate Buffer (pH 6.0; Sigma, C9999) for antigen retrieval. Slides were blocked with 3% Bovine serum albumin (Sigma #A2153) for 1 hour at room temperature and incubated overnight in primary antibody at 4°C. Slides were visualized with fluorophore-conjugated secondary antibody. For Picrosirius Red stainings, deparaffinized FFPE sections were incubated for 2 hours in a solution consisting of 0.01% fast green (Sigma, F7258-25G) in picric acid (Sigma, P6744-1GA), and then incubated for an additional 2 hours in a solution consisting of 0.04% fast green, 0.1% Sirius Red (Sigma, 365548-25G) in picric acid. Slides were then rehydrated and mounted in Permount solution (Fisher SP15) for microscopic evaluation. For immunostaining on frozen sections, freshly collected liver tissue was embedded directly into OCT blocks on dry ice and stored at -80°C. To make cryosections tissue embedded in OCT blocks were sectioned using cryostat and mounted onto charged glass slides. Cryosections were fixed with -20°C Methacarn (60% Methanol, 30% Chloroform, 10% Acetic Acid) for 10 minutes at -20°C. Slides were then washed and incubated for 1 hour at 37C in primary antibody. Fluorophore-conjugated secondary antibody was used for visualization. Slides were mounted using fluoromount with DAPI (Southern Biotech #0100-20).

### Digital Pathology Analysis by Pharmanest

Picrosirius Red stained liver sections were imaged at 40X magnification using a whole slide scanner (Hamamatsu NanoZoomer) with tiling across the entire slide. Images were analyzed by a commercial partner (Pharmanest, Princeton, NJ) using a single-fiber, high-content image analysis platform that assess collagen deposition, fiber morphometry, and fibrosis architecture to generate a Phenotypic Fibrosis Score (Ph-FCS), which is a continuous, quantitative measure of fibrosis severity reflecting fibrosis severity and based on calibrated metrics of fiber length, density, and orientation.

### Single Nucleus Isolation and Sequencing

For snRNA-seq of mouse samples, 40 to 60 mg of total liver tissue (pooled from 3 mice per condition) were chopped on ice in 1 mL of 0.03% TST buffer [146 mM NaCl, 10 mM Tris-HCl (pH 7.5), 1 mM CaCl_2_, and 21 mM MgCl_2_ with freshly added 0.03% Tween 20, 0.01% BSA, and RiboLock RNase Inhibitor (0.4 U/mL; Thermo Fisher Scientific, FEREO0382) (30)] with Tungsten Noyes Spring Scissors (FST, 15514-12). The resulting nuclei suspension was then filtered through 40-μm cell strainers (Thermo Fisher Scientific, 352340) into a fresh 50-mL conical tube on ice.

One more milliliter of 0.03% TST was used to rinse the cell strainer, and 3 mL of ST buffer were added to the nuclei suspension, followed by briefly mixing by gentle flicking. The nuclei suspensions were spun at 500xg for 5 min at 4°C in a swing-bucket centrifuge. The final nuclei pellet was suspended in 200 μL of 0.4% BSA in PBS by pipetting 30 times with a regular p1000 tip. The resulting pellet was resuspended by pipetting 30 times in 200 µL of 0.4% BSA/PBS, and nuclei were visually inspected using a 1:1 trypan blue solution under a light microscope. Samples were transferred to the Mount Sinai Genomics CoRE for counting, quality control and sequencing. Nuclei preparations were processed by the Chromium 3′ Gene Expression V2 Kit according to the manufacturer’s guidelines. Sequenced files from each independent sample were processed through 10X Genomics Space Ranger software to generate a count matrix.

### Single Cell Isolation and Sequencing

For scRNA-seq, 40mg of liver tissue each from 5 mice were pooled and stored in RPMI (Corning, 15-040-CM) + 10% fetal bovine serum (FBS; Invitrogen, 10082147) on ice overnight. The following morning, livers were injected with digestion buffer consisting of RPMI with 10% FBS, 0.25mL Collagenase IV (Sigma, C5138), and 0.1mg/mL DNase I (Sigma, D5025). For digestion, samples were initially minced and incubated at 37°C for 1 hour under slow agitation (80rpm), and then mechanically digested using 16G needles, before being filtered through 100uM and 70uM strainers. Thereafter, for gradient separation, samples were centrifuged (500xg, 4°C, 10 min), resuspended in 25% Percoll, and layered atop 70% Percoll (Cytiva, 17-0891-02), and centrifuged again (500g, 20°C, 25 min, no acceleration/brake). Non-parenchymal cells (NPCs) were collected at the meeting point of the Percoll layers. NPCs were rinsed with 1XHBSS (no ions; Corning, 21-022-CV) and subjected to RBC lysis (Fisher, A1049201). A fraction of cells from this suspension was transferred to the Mount Sinai Genomics CoRE for counting, quality control and sequencing. Nuclei preparations were processed by the Chromium 3′ Single Cell 3’ Reagent Kit according to the manufacturer’s guidelines. 10X Genomics Cell Ranger software v6 was used to generate a count matrix from the sequenced files from each sample.

### Cell Abundance, Velocity, and Pathway Analyses

Single-cell and single-nucleus RNA sequencing data processed by Cell Ranger were analyzed with Seurat [26] (v5; using default parameters). After importing the Cell Ranger output count matrix, ambient RNA was first removed using Cellbender [27] (v0.3.2; using default parameters). Cells with a sufficient number of detected genes (more than 200 but fewer than 2500) and less than 20% mitochondrial gene content were retained. QC metrics were iteratively refined and processed with violin and scatter plots. Integration across all time points (peak, 2-week regression, and 6-week regression) was performed using Harmony [28] (v0.1) to correct for batch effects and ensure that data from all time points could be directly compared. Clustering was done with Seurat (default parameters), and clusters were annotated by examining top marker genes for each group. Abundance of each cell cluster at different time points was compared. Spliced and unspliced mRNA counts were exported and integrated into scVelo’s analysis pipeline. Using scVelo [29] (v0.3.3 ; default parameters), velocity vectors across all cells were generated and overlaid onto the UMAP embeddings generated from Seurat integration. For pathway analysis, genes in each cluster exhibiting a log_2_fold change greater than 1 compared to other clusters were inputted into AmiGO2 [30], a web-based Gene Ontology (GO) tool, to perform functional enrichment analysis (Statistical enrichment test, Mann-Whitney U test, two-sided; p<0.05).

### Deconvolution and Integration of Spatial RNA Sequencing Data

To increase resolution and sensitivity of spatial transcriptomics data, we integrated snRNA-seq data with spatial Visium datasets using Tangram [31] (v1.0.4). For each spatial sample, Visium data were read and preprocessed using Scanpy, by filtering out lowly expressed genes (present in fewer than three cells), total-count normalization to a target sum of 10,000 counts per spot, log transformation, and the selection of 2,000 highly variable genes. Tangram was applied in “cells” mode with a density prior based on RNA counts, and mapping was run for 500 epochs on a GPU-enabled device.

### Spatial Sequencing

For spatial sequencing, two complementary experiments were performed using liver samples from the Peak CCl_4_ Fibrosis and 6-Week CCl_4_ Regression conditions. In the first experiment, cryosections were prepared from liver samples that were harvested and stored directly on OCT. Two sections were placed on each slide and samples were sent to the Mount Sinai Genomics CoRE for library preparation and sequencing based on the standard 10X Visium protocol for fresh frozen tissue. In the second experiment, two sections of formalin-fixed, paraffin-embedded (FFPE) liver blocks were placed on each slide and sent to the Mount Sinai Genomics CoRE for processing. To generate a count matrix, the sequenced files from each sample were processed through 10X Genomics Space Ranger software.

### Cell-cell Interaction Predictions

Cell–cell communication based on sc/snRNAseq data was predicted using 3 different software methods. CellphoneDB [17] (v5) was used with default parameters to identify statistically significant ligand–receptor interactions between clusters in our sc/snRNAseq datasets. In parallel, a previously established method called “Calligraphy” [19] was used to construct a detailed, directed network of gene modules based on these interactions. Gene modules were identified through correlation analysis and refined using community detection algorithms as defined [19]. Each module, which comprises a set of co-expressed genes, was represented as a node in a network graph. Directed edges between nodes were established by quantifying the number of known ligand–receptor interactions connecting genes from one module to another.

### *In vivo* rSFRP2 Treatment

To induce MASH, 6-week-old C57B/6J male mice (Jackson Laboratory) were placed on the FAT-MASH protocol for 6 weeks [15]. Some of these mice, designated as “Peak MASH Fibrosis”, were euthanized 48 hours after the final CCl_4_ injection to assess maximum fibrosis response. In separate groups, mice were allowed to recover for two and six weeks with twice weekly treatment with rSFRP2 (200ng/treatment, diluted in PBS; R&D, 1169-FR-025/CF) (designated as “2wk SFRP2 Regression” and “6wk SFRP2 Regression”) or with PBS vehicle control (Gibco, 10010-023) (designated as “2wk Saline Regression” and “6wk Saline Regression”).

### *In situ* zymography with activatable zymography probes

Staining with Activable Zymography Probes (AZPs) was performed as described previously1,2. Briefly, tissue sections were air-dried, fixed in pre-chilled acetone (−20 °C) for 10 minutes, and then air-dried again. Slides were rehydrated in phosphate-buffered saline (PBS) with three 5-minute washes. Samples were then incubated at room temperature for 30 minutes in a protease assay blocking buffer consisting of 50 mM Tris-HCl, 300 mM NaCl, 10 mM CaCl_2_, 2 mM ZnCl_2_, 0.02% (v/v) Brij-35, and 1% (w/v) bovine serum albumin (BSA), adjusted to pH 7.4. After removing the blocking buffer, a staining solution containing fluorescently conjugated activity-based probes (AZPs, 1 μM each) along with a 0.1 μM poly-arginine (polyR) control was applied. Slides were incubated at 37 °C in a humidified chamber for 4 hours. In co-staining experiments, primary antibodies were included directly in the AZP incubation solution. These included Desmin (ab227651, Abcam), CK7 (ab181598, Abcam), Collagen Type I (600-401-103S, Rockland), and von Willebrand Factor (ab11713, Abcam). After incubation with probes, slides were washed three times in PBS (5 minutes each), counterstained with Hoechst 33342 (5 μg/mL, Invitrogen), and labeled with fluorescent secondary antibodies. Final washes in PBS were followed by mounting using ProLong Diamond Antifade Mountant (Invitrogen). Imaging was performed using a Pannoramic 250 Flash III slide scanner (3DHistech). For protease inhibition studies, AEBSF (400 μM), marimastat (1 mM), or a protease inhibitor cocktail (P8340, Sigma Aldrich) supplemented with AEBSF and marimastat was added during both the blocking and probe incubation steps.

## Statistical analysis

Results are shown as means ± SEM unless indicated otherwise. Statistical analysis was performed using Prism unless otherwise specified. Student’s t test was used when comparing two groups with similar variances (F test performed to compare variance, P > 0.05). Analysis involving multiple groups was performed using one-way or two-way analysis of variance (ANOVA) followed by post hoc t test. “*” = p <0.05, “**” = p < 0.01, “***” = p < 0.001, “****” = p < 0.0001.

