## Supplemental figures and table for "Scar-associated endothelial-stellate cellular crosstalk drives fibrosis resolution in MASH"

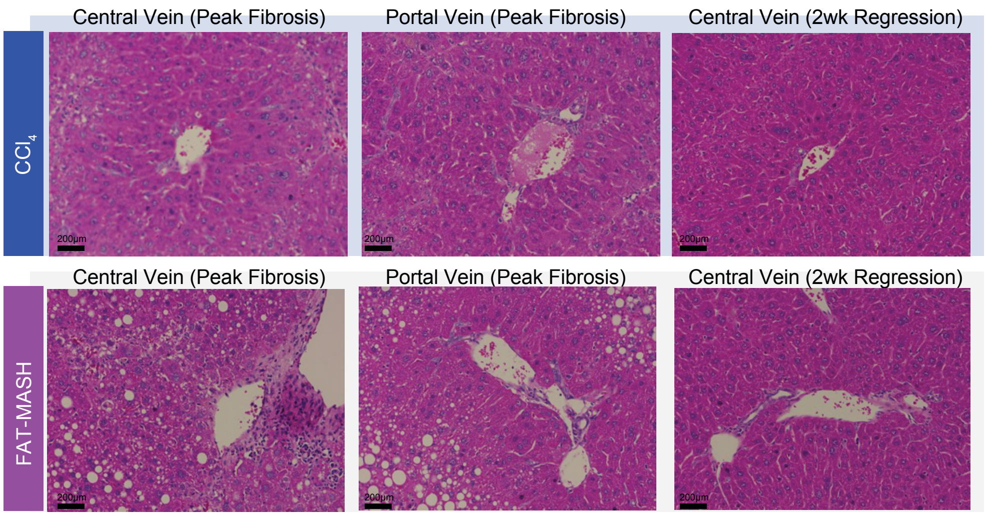


**Figure S1**. Steatosis and inflammation resolve in MASH model at 2wks Regression.

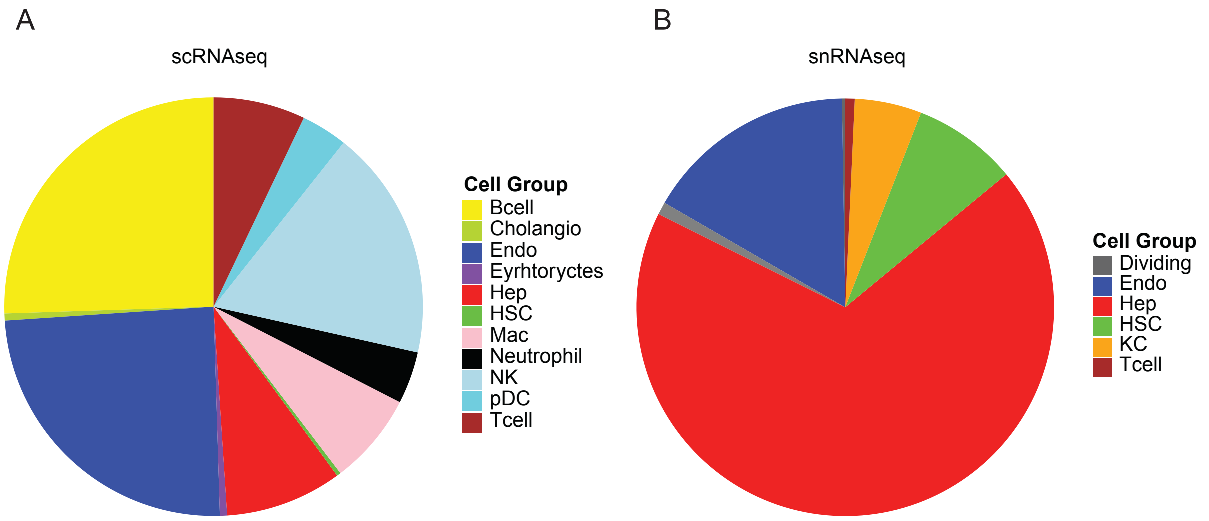


**Figure S2**. Proportional cell type capture by scRNAseq (A) and snRNAseq (B).

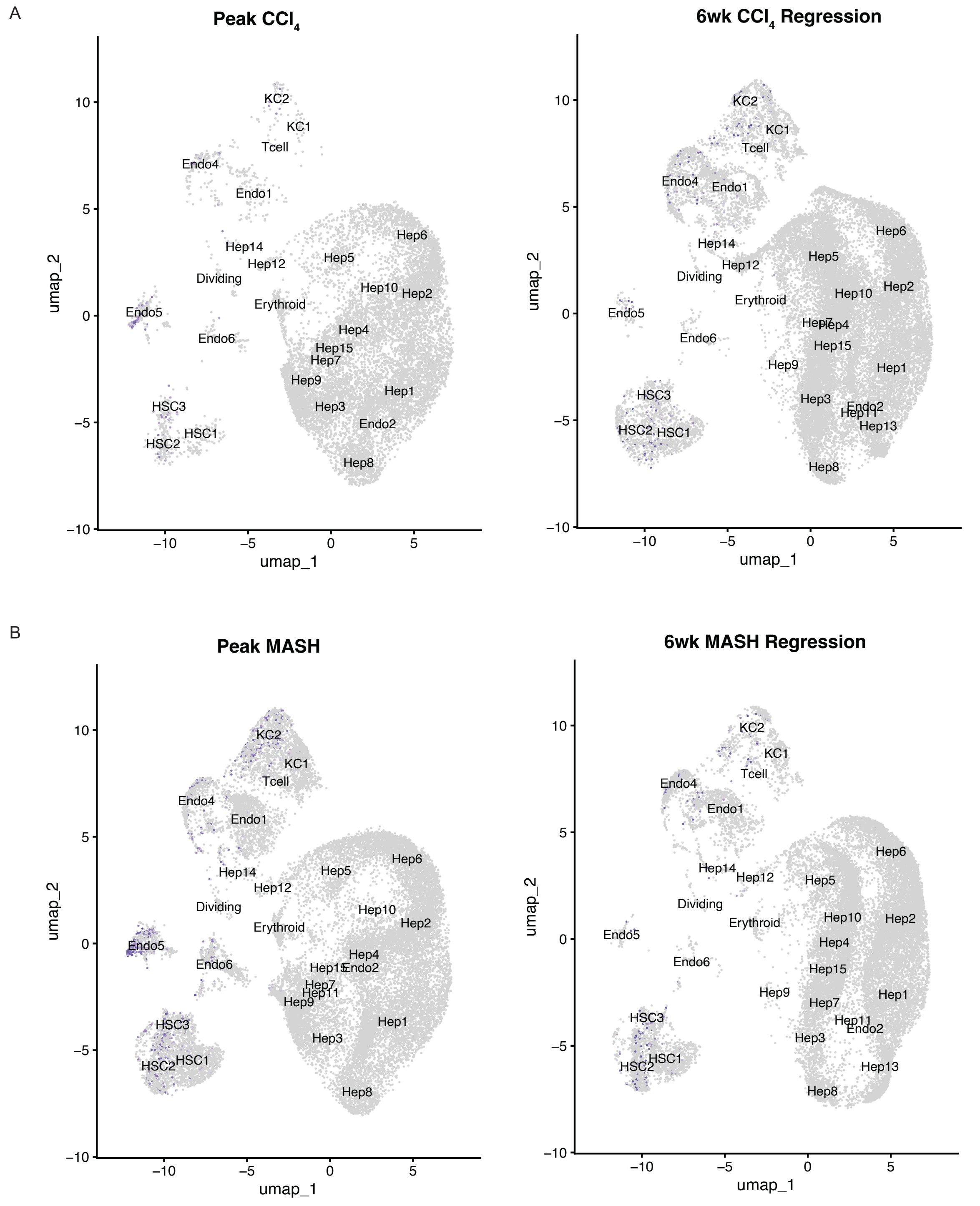


**Figure S3**. Vimentin expression in a subset of endothelial cells that disappears at 6wks Regression.

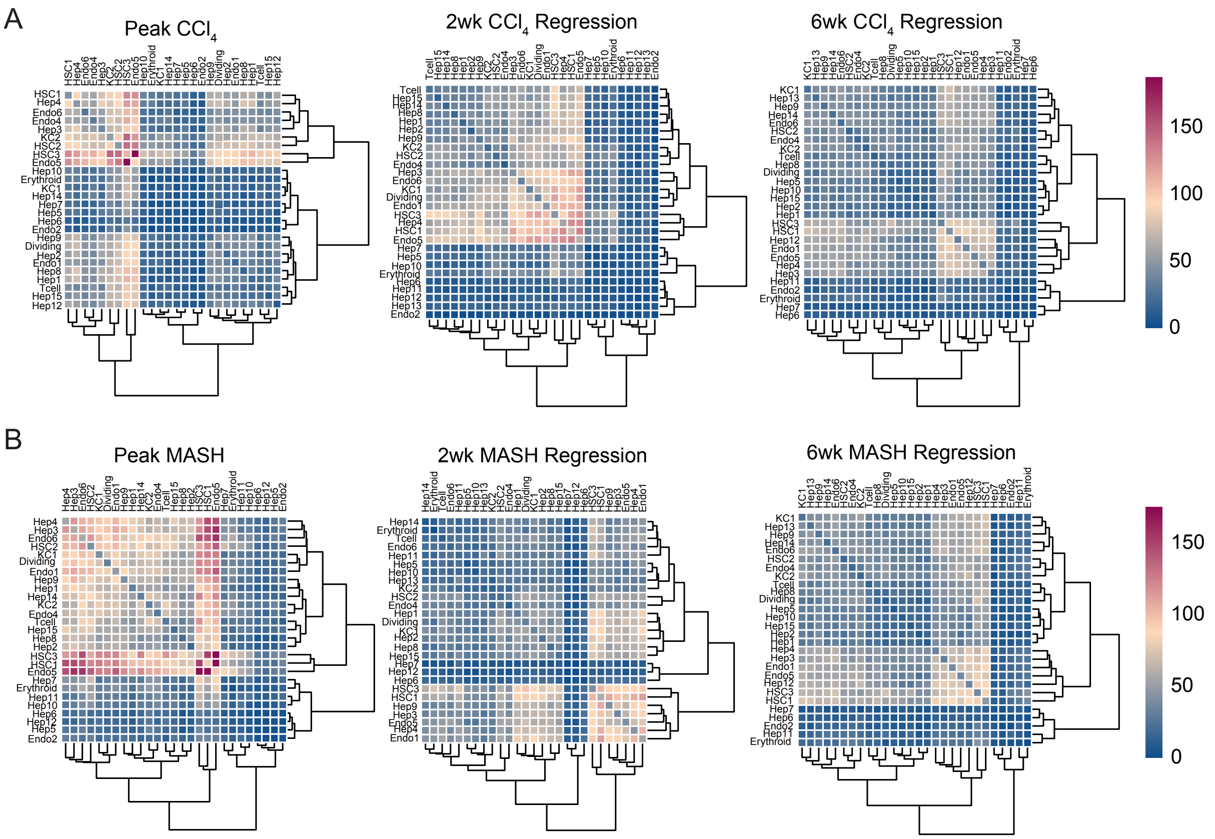


**Figure S4**. Global reduction in cell-cell communication with fibrosis resolution. CellphoneDB analysis of $CCl_4$ (A) and MASH Regression (B) snRNAseq datasets. Heatmaps show the absolute number of receptor-ligand interactions between each cell type pair at each of the time points.

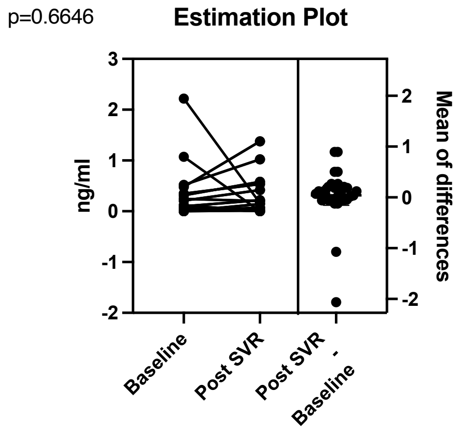


**Figure S5**. SFRP2 levels in paired HCV patient serum before and post-SVR. Serum SFRP2 levels in patients measured by ELISA, p-value calculated by paired-t test.

A

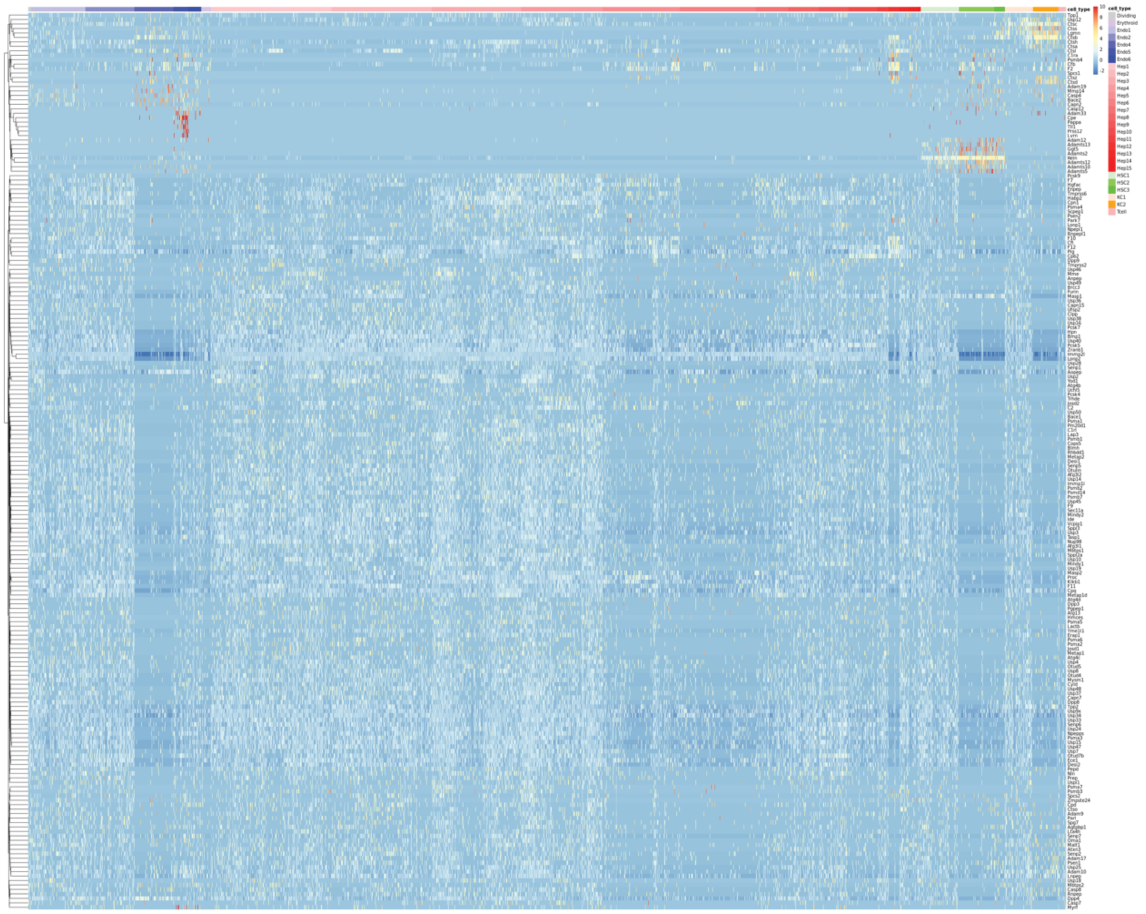
cell_type
10
8
6
4
2
0
-2
cell_type
Dividing
Erythroid
Endo1
Endo2
Endo4
Endo5
Endo6
Hep1
Hep2
Hep3
Hep4
Hep5
Hep6
Hep7
Hep8
Hep9
Hep10
Hep11
Hep12
Hep13
Hep14
Hep15
HSC1
HSC2
HSC3
KC1
KC2
Tcell

B

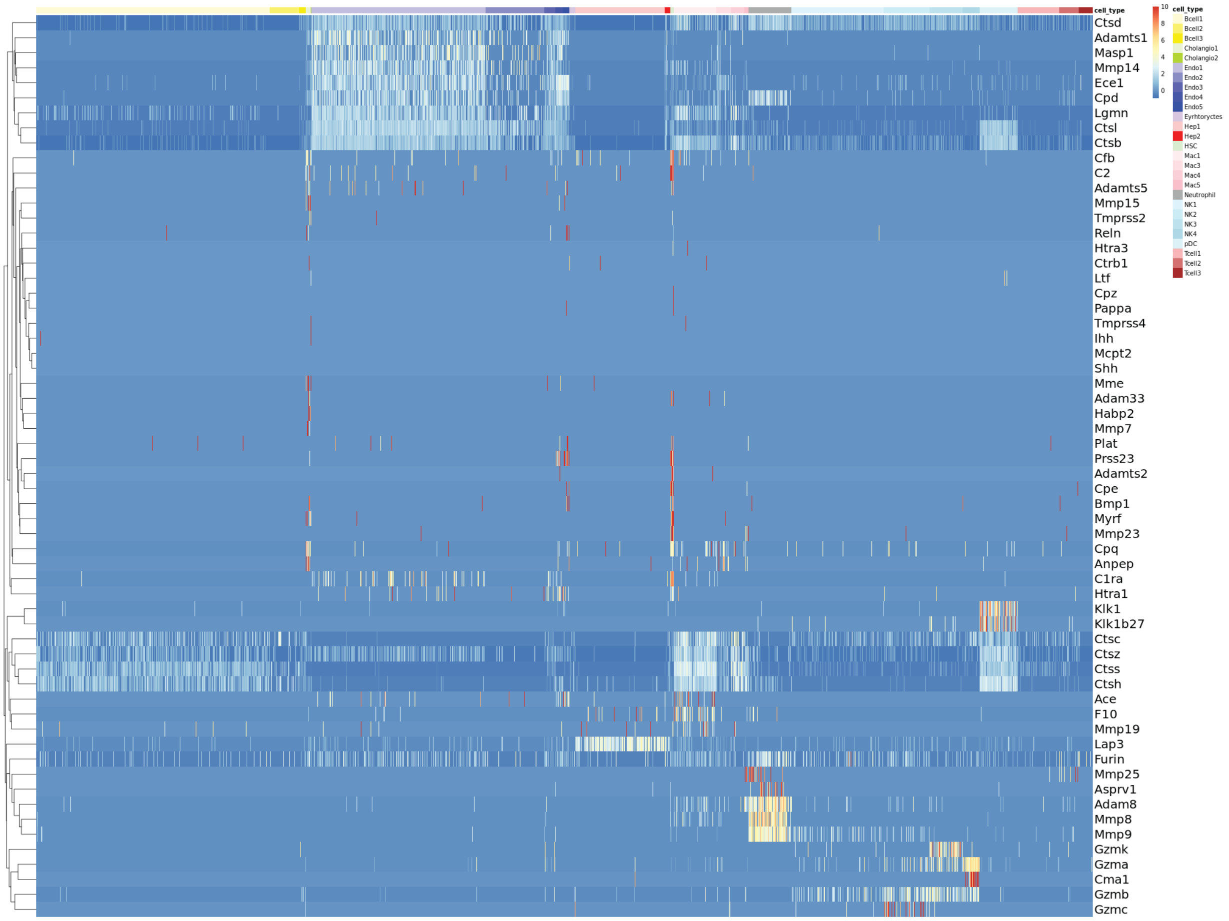


**Figure S6**. Landscape of protease expression in MASH and $CCl_4$ Regression mouse models. Heatmap showing protease expression in each cell cluster from the integrated (**A**) single nucleus and (**B**) single cell sequencing data of the CCl4-only and MASH Regression model mice.

**Table S1**: Cell-cell interactions predicted by Calligraphy. Significant receptor-ligand module interactions identified by Calligraphy in the integrated MASH and $CCl_4$ snRNAseq datasets.

| Peak Fibrosis | | | 2wk Regression | | | 6wk Regression | | |
|---|---|---|---|---|---|---|---|---|
| **Cell Type 1** | **Cell Type 2** | **Interaction** | **Cell Type 1** | **Cell Type 2** | **Interaction** | **Cell Type 1** | **Cell Type 2** | **Interaction** |
| Neutrophil | HepA | Igf1r – Igf1 | HSCA | CholangioB | Sfrp1 – Wnt7a | BcellA | NK/BcellB | Cd22 – Ptprc |
| Neutrophil | HepA | Fpr2 – Saa1 | HSCA | CholangioB | Sfrp1 – Wnt7b | Neutrophil | NK/BcellA | Ccr1 – Ccl3 |
| CholangioB | CholangioA | Dsg2 – Dsc2 | HSCA | CholangioB | Sfrp2 – Wnt7a | Neutrophil | NK/BcellA | Ccr1 – Ccl5 |
| CholangioB | HSCA | Epha7 – Efna5 | HSCA | CholangioB | Sfrp2 – Wnt7b | Neutrophil | HepA | Igf1r – Igf1 |
| CholangioB | HSCA | Erbb4 – Ereg | HSCA | Endo4 | Sfrp1 – Wnt9b | | | |
| HSCA | CholangioA | Sfrp2 – Wnt7b | HSCA | Endo4 | Sfrp2 – Wnt9b | | | |
| BcellA | NK/BcellA | Cd22 – Ptprc | Neutrophil | NK/BcellA | Ccr1 – Ccl3 | | | |
| Neutrophil | NK/BcellA | Ccr1 – Ccl3 | Neutrophil | NK/BcellA | Ccr1 – Ccl5 | | | |
| Neutrophil | NK/BcellA | Ccr1 – Ccl5 | CholangioB | HSCA | Fgfr2 – Fgf2 | | | |
| CholangioA | EndoA | Pigr – Aplp2 | CholangioB | HSCA | Fgfr2 – Fgf7 | | | |
